# Chronic fluoxetine treatment desensitizes serotoninergic inhibition of GABA inputs and the intrinsic excitability of dorsal raphe serotonin neurons

**DOI:** 10.1101/2024.05.07.592963

**Authors:** Wei Zhang, Ying Jin, Fu-Ming Zhou

**Author notes:** Corresponding author: Fu-Ming Zhou, Department of Pharmacology, University of Tennessee College of Medicine, Memphis, TN 38103,.

## Abstract

Dorsal raphe serotonin (5-hydroxytryptamine, 5-HT) neurons are spontaneously active and release 5-HT that is critical to normal brain function such mood and emotion. Serotonin reuptake inhibitors (SSRIs) increase the synaptic and extracellular 5-HT level and are effective in treating depression. Treatment of two weeks or longer is often required for SSRIs to exert clinical benefits. The cellular mechanism underlying this delay was not fully understood. Here we show that the GABAergic inputs inhibit the spike firing of raphe 5-HT neurons; this GABAergic regulation was reduced by 5-HT, which was prevented by G-protein-activated inwardly rectifying potassium (Girk) channel inhibitor tertiapin-Q, indicating a contribution of 5-HT activation of Girk channels in GABAergic presynaptic axon terminals. Equally important, after 14 days of treatment of fluoxetine, a widely used SSRI type antidepressant, this 5-HT inhibition of GABAergic inputs was substantially downregulated. Furthermore, the chronic fluoxetine treatment substantially downregulated the 5-HT activation of the inhibitory Girk current in 5-HT neurons. Taken together, our results suggest that chronic fluoxetine administration, by blocking 5-HT reuptake and hence increasing the extracellular 5-HT level, can downregulate the function of 5-HT1B receptors on the GABAergic afferent axon terminals synapsing onto 5-HT neurons, allowing extrinsic, behaviorally important GABA neurons to more effectively influence 5-HT neurons; simultaneously, chronic fluoxetine treatment also downregulate somatic 5-HT autoreceptor-activated Girk channel-mediated hyperpolarization and decrease in input resistance and intrinsic excitability, rendering 5-HT neurons resistant to autoinhibition and leading to increased 5-HT neuron activity, potentially contributing to the antidepressant effect of SSRIs.

## Introduction

The serotonin (5-hydroxytryptamine, 5-HT) neurons in the dorsal raphe nucleus (DRN) project to every forebrain areas in rodents and primates including humans (Baker et al. 1990; Charara and Parent 1998; Commons 2016; Hornung 2003; Parent et al. 2011; Smiley & Goldman-Rakic 1996; Steinbusch 1981). 5-HT, the key neurotransmitter of these neurons, contributes critically to the development of these brain areas and also to the normal function of the brain in adult (Mosienko et al. 2015; Teissier et al. 2017), as indicated by the fact that 5-HT agonists such as lysergic acid diethylamide (LSD) can alter human cognition and triggers powerful hallucination (Kraehenmann et al. 2017; Preller et al. 2018). Further supporting the importance of the 5-HT system, selective serotonin reuptake inhibitors (SSRIs) that increases the extracellular 5-HT level are effective for treating depression symptoms (Wong et al. 2005). Thus, controlling and regulating 5-HT neuron spiking activity that trigger 5-HT release in the target area is important for normal brain function and may contribute to the pathogenesis of depression and other neuropsychiatric-behavioral disorders.

5-HT neurons often fire around 3 Hz in a quiet awake state (Dougalis et al. 2017; Jacobs and Azmitia 1992; Jacobs et al. 2002; Sakai 2011; Tuckwell and Penington 2014). This tonic firing is altered when the animal’s mental and behavior state and its environment are changed. Synaptic inputs likely contribute to the behavior-related changes in 5-HT neuron firing. Studies have indicated that GABA inputs inhibit raphe 5-HT neuron firing (Levine and Jacobs 1992; Wang and Aghajanian 1977). For example, the brain 5-HT system/5-HT neurons may contribute critically to the induction and regulation of sleep; 5-HT neuron destruction can lead to insomnia (Jouvet 1999); and REM sleep is associated with inhibition of 5-HT neuron spike firing or decreased 5-HT release (Iwasaki et al. 2018; Jouvet 1999; Nitz and Siegel 1997; Sakai and Crochet 2001; Siegel 2004), or 5-HT neuron activity may inhibit REM sleep (Saper et al. 2010)--although this is not fully settled (Sakai 2011). This inhibition of 5-HT neuron activity may be mediated, at least partially, by GABAergic input from extrinsic sources such as the hypothalamus, lateral habenula nucleus, mesopontine rostromedial tegmental nucleus (RMTg), lateral preoptic area and the pontine ventral periaqueductal gray including the DRN, VTA, SNr (Bernard et al. 2012; Gervasoni et al. 2000; Kirouac et al. 2004; Lavezzi et al. 2012; Pollak et al. 2014; Reisine et al. 1982; Sego et al. 2014; Soiza-Reilly and Commons 2014; Taylor et al. 2014; Wang and Aghajanian 1977; Zhou et al. 2017). These GABAergic inputs can affect the spike firing of the raphe 5-HT neurons, which in turn affects the synaptic 5-HT level in 5-HT neuron projection areas and hence behaviors. Thus, this GABAergic input is important. The GABAergic afferents may express 5-HT1B receptors that may inhibit GABA release (Lemos et al. 2006).

In addition to GABAergic inhibition by other neurons, raphe 5-HT neurons can also inhibit themselves (autoinhibition) by expressing inhibitory 5-HT1A autoreceptors (Descarries & Riad 2013; Stamford et al. 2000). 5-HT1A receptor activation triggers, via Gi/o beta-gamma subunits, the opening of G-protein activated inwardly rectifying K (Girk) channels that are highly expressed in raphe 5-HT neurons, leading to a hyperpolarization and inhibition of these neurons (Bayliss et al. 1997; Lei et al. 2000; Dougalis et al. 2017; Penington et al. 1993a,b; Montalbano et al. 2015; Saenz del Burgo et al. 2008; Tuckwell and Penington 2014). Furthermore, chronic (≥ 2 weeks) fluoxetine treatment, matching the required duration for clinical antidepressant effects to appear, has been shown to downregulate or desensitize 5-HT1A receptors (Blier and El Manari 2013; Descarries and Riad 2013; Hensler 2002; Rainer et al. 2012) and remove the initial 5-HT neuron firing inhibition after selective 5-HT reuptake inhibitor treatment, detected by in vivo extracellular spike (Blier and de Montigny 1983; Czachura and Rasmussen 2000; Guiard et al. 2012). This 5-HT1A receptor downregulation has been suggested to be critical to fluoxetine’s antidepressant effect (Wong et al. 2005). Indeed, molecular genetics-mediated inactivation or reduction of these 5-HT1A autoreceptors has been reported to have antidepressant-like effects (Richardson et al. 2010; Ferrés-Coy et al. 2013). In our present study, we sought to complement these published findings by chronically treating mice with fluoxetine and then performing whole-cell patch-clamp recording to provide more direct and detailed information about the potential effect of chronic fluoxetine on 5-HT-activated, Girk-mediated hyperpolarization and the associated changes in intrinsic excitability in dorsal raphe 5-HT neurons.

## Materials and Methods

### Animals

All experimental protocols were approved by the Institutional Animal Care and Use Committee at University of Tennessee Health Science Center, Memphis, Tennessee (animal protocol # 17-033.0). Breeder mice (C57Black/6J) were purchased from The Jackson Laboratory. Mice had free access to food and water. The room light was on 7:00AM to 7:00 PM and off for the night. Male and female mice (starting on PN 13 days) were administered intraperitoneally the antidepressant fluoxetine (10 mg/kg) or saline for two weeks (once daily). The 2-week duration was mainly based on the published results of Blier and de Montigny (1983) and Czachura and Rasmussen (2000). Fluoxetine was chosen because it is a commonly prescribed and hence important antidepressant. The fluoxetine dose (10 mg/kg) was based on the results of Czachura and Rasmussen (2000).

### Brain slice preparation

Wild-type male and female mice (PN27-30) were rapidly decapitated and their brains were quickly dissected out. The brainstem was rapidly removed and placed in ice-cold, high-sucrose, artificial cerebral spinal fluid (ACSF) that contained the following (in mM): 220 Sucrose, 2.5 KCl, 1.25 NaH_2_PO_4_, 2 ascorbic acid, 25 NaH_2_CO_3_, 0.5 CaCl_2_, 7 MgCl_2_, 20 D-glucose, pH 7.4 (continuously bubbled with 95% O_2_-5% CO_2_). Three-four coronal midbrain slices containing the dorsal raphe nucleus were cut (300 μm thick) using a Leica vibratome (VT1200S, Leica Microsystems, Germany) and immediately transferred to a holding chamber filled with the normal extracellular solution (in mM): 2 ascorbic acid, 125 NaCl, 2.5 KCl, 1.25 NaH_2_PO_4_, 25 NaH_2_CO_3_, 2.5 CaCl_2_, 1.3 MgCl_2_, 10 D-glucose, pH 7.4 (when continuously bubbled with 95% O_2_-5% CO_2_)--this extracellular solution was also the normal perfusing solution. After incubation at 34 °C for 30 minutes, slice were maintained at room temperature (22-24 °C) until being transferred to the recording chamber on the stage of the microscope. Drugs were applied via the perfusing solution. The perfusion rate was 2 ml/min.

### Electrophysiological recording

Conventional whole-cell patch-clamp recordings were performed in the recording chamber maintained at 30 °C. As illustrated in **Fig. 1A,B**, the dorsal raphe neurons were visualized in the midline region ventral to the aqueduct, using an Olympus upright microscope (BX51WI) fitted with a 60×water-immersion lens and DIC optics and a Zeiss Axiocam MRm digital camera. The slice was submerged in and perfused with normal extracellular solution equilibrated with 95% O_2_-5% CO_2_. Whole-cell recording pipettes were pulled from borosilicate glass capillary tubing (KG-33, 1.65 mm outer diameter, 1.10 mm inner diameter, King Precision Glass, Claremont, CA) on a two-stage PC-10 micropipette puller (Narishige, Tokyo, Japan).

**Fig. 1.**
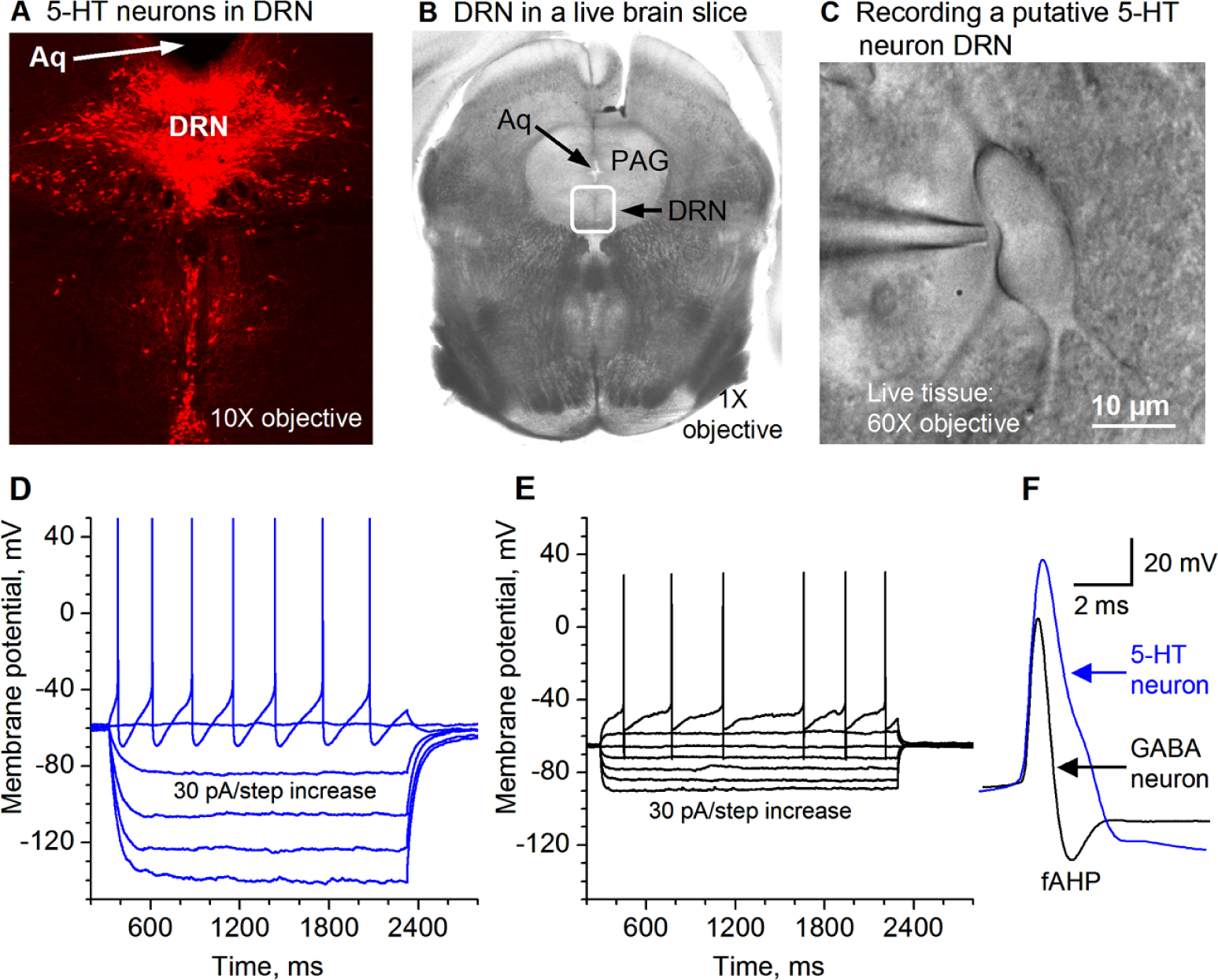
Recording and identification of 5-HT neurons in the dorsal raphe nucleus (DRN) in a mouse. **A.** Tryptophan hydoxylase (TPH) immunostain identifies 5-HT neurons in mouse DRN. Aq, aqueduct. **B.** A live coronal brain slice containing DRN viewed under a low power objective. The boxed area is roughly the DRN. Aq, aqueduct. PAG: periaqueductal gray. **C.** A putative 5-HT neuron in DRN being patch-clamped. **D.** Membrane potential responses to current injection in example putative 5–HT neuron in DRN. The current strength started at −120 pA and the step size was 30 pA. **E.** Membrane responses to current injection in example putative GABA neuron in DRN. The current strength started at −120 pA and the step size was 30 pA. **F.** High temporal resolution display showing that action potential duration and amplitude for DRN 5-HT neurons are clearly longer and larger than those for DRN GABA neurons.

A potassium methanesulfonate (KSO_3_CH_3_)-based low Cl^-^ intracellular solution (in mM: 135 KSO_3_CH_3_, 5 KCl, 0.5 EGTA, 10 HEPES, 2 Mg-ATP, 0.2 Na-GTP, and 4 Na_2_-phosphocreatine, pH 7.25, 280-290 mOsm) was used for recording the natural hyperpolarizing GABA_A_ inhibitory postsynaptic potentials (IPSPs) and their effects on spiking activity (**Fig. 2**). A high Cl^-^ intracellular solution (in mM: 135 KCl, 0.5 EGTA, 10 HEPES, 2 Mg-ATP, 0.2 Na-GTP, and 4 Na_2_-phosphocreatine, pH 7.25, 285 mOsm) was used for recording inward GABA_A_ inhibitory postsynaptic currents (IPSCs) at –70 mV. The patch pipette resistance was about 2 MΩ. Access resistance change was <15% during recording. Liquid junction potentials were not corrected.

Evoked IPSCs (eIPSC) were electrically stimulated via a bipolar stimulating electrode; stimulus timing and intensity was controlled by a Master-8 pulse generator (A.M.P.I.) and an A360 stimulus isolator (World Precision Instruments, Sarasota, FL), in the presence of ionotropic glutamate receptor blockers 6-cyano-7-nitroquinoxaline-2,3-dione (CNQX, 10 μM, to block non-NMDA glutamate receptors) and D-(−2-amino-5-phosphonopetanoic acid (D-APV, 20 μM, to block NMDA receptors). The dorsal raphe neuron under recording was usually 100 μm away from the two tips of stimulating electrode. Stimulus intervals were 30 s unless otherwise stated. Input resistance was measured in the whole cell configuration in current-clamp mode by injecting current pulse of –20 pA with a duration of 1000 ms.

### Data acquisition and analysis

Cells were recorded in the whole cell configuration in voltage- and current-clamp. Recordings were made with an Axopatch 200B patch-clamp amplifier (Axon Instruments, USA) and pCLAMP 9.2 software (Clampex for data acquisition and Clampfit for data analysis). Signals were filtered at 5 kHz, digitized at 10 kHz with the Digidiata 1322A interface (Axon Instruments), and stored in a computer hard disk for off-line analysis. Data are presented as mean ± SEM. ANOVA and paired and unpaired student’s t-tests were used for statistical comparison. *p* < 0.05 were considered significant.

### Drugs

Picrotoxin, DL-2-amino-5-phosphonopentanoic acid (AP-5), 6-cyano-7-nitroquinoxaline-2,3-dione (CNQX), 5-HT, tetrodotoxin (TTX), tertiapin-Q, were purchased from Tocris or Sigma. Fluoxetine was either purchased from Tocris or supplied by NIMH Drug Supply program. 5-HT was purchased from Sigma, prepared as a 10 mM stock solution in pure water, and stored at - 20 °C and was used at the final concentration of 10 μM.

### DRN 5-HT neuron mapping with tryptophan hydoxylase (TPH) immunostain

To guide our electrophysiological recording, we also immunostained the tissue sections for TPH, a key 5-HT synthesis enzyme, to visualize DRN 5-HT neurons (**Fig. 1A**). We used conventional immunohistochemical procedures we have published (Ding et al. 2015a,b; Li et al. 2015; Zhou et al. 2009). Briefly, free-floating sections (50 μm in thickness) were incubated with 2% fat-free milk, 1% bovine serum albumin (BSA), and 0.8% Triton X-100 in 0.01 M PBS for 1 hour at room temperature to block nonspecific staining. Then these free-floating tissue sections were incubated with a primary mouse anti-TPH antibody (diluted 1:500, purchased Sigma) for 48 hours at 4 °C. After 3 × 10 min PBS rinses to wash out excess primary antibody, the tissue sections were incubated with Alexa Fluor 568 (red) donkey anti-mouse secondary antibody (diluted 1:200, purchased from Invitrogen) for 2 hours at room temperature, followed by rinsing 3 × 10 min, cover-slipped, and digitally photographed on a fluorescence microscope.

## Results

### Electrophysiological identification of dorsal raphe 5-HT neurons

Under visual guidance of video-microscope equipped with a 60X water immersion lens and DIC optics, we made whole-cell patch clamp recordings in neurons in the dorsal raphe of mice. To record putative 5-HT neurons, we targeted relatively large neurons in DRN (∼ 20-25 μm in the longest dimension, **Fig. 1B,C**) because literature data indicate that large neurons in the dorsal raphe are 5-HT neurons. Our presumed 5-HT neurons had the following basic electrophysiological properties: resting membrane potential of –61.3 ± 1.2 mV (n=18 cells), high spike peak amplitude of 93.3 ± 1.3 mV (n=18), a wide action potential duration of 2.94± 0.11 m (n=18) at the spike base and a whole-cell input resistance of 362.2 ± 5.4 MΩ (n=18). These basic intrinsic membrane properties are consistent with those reported for dorsal raphe 5-HT neurons in the literature (Liu et al. 2002). Although not our focus in this study, for a purpose of comparison, we also recorded 10 presumed GABA neurons by targeting smaller neurons (≤15 μm in the longest dimension). As shown in **Fig. 1D,E,F**, these presumed GABA neurons had the following electrophysiological properties that are clearly different from those in the presumed 5-HT neurons: action potential duration was 1.19 ± 0.11 ms at the base (n=10 cells), action potential amplitude was 76.4 ± 2.3 mV, and the input resistance was 185.4±8.5 MΩ (n=10 cells); these putative GABA neurons also had a deep fast afterhyperpolarization (fAHP) (**Fig. 1D,E,F**). Equally important, our presumed 5-HT neurons were hyperpolarized by 5-HT. These characteristics are consistent with electrophysiological parameters for dorsal raphe 5-HT and GABA neurons in the literature (Allers and Sharp 2003; Haj-Dahmane 2001; Liu et al. 2002, 2005; Mlinar et al. 2016), indicating that under our recording conditions, 5-HT neurons can be reasonably identified based on the whole-cell electrophysiological properties.

### GABAergic inhibition of DR 5-HT cell firing

We first determine if dorsal raphe 5-HT neurons receive IPSPs and how these IPSPs affect the spiking activity, under the condition with fast ionotropic glutamate receptors were blocked by 10 μM CNQX and 20 μM D-APV. As shown in **Fig. 2**, local electrical stimulation evoked typical hyperpolarizing IPSPs and these IPSPs clearly and obviously inhibited the generation of spontaneous action potentials in DR 5-HT neurons (n=6). These data indicate that regulation of the GABAergic inputs may exert an important influence on 5-HT neuron activity, providing the functional importance for the experiments described below.

**Fig. 2.**
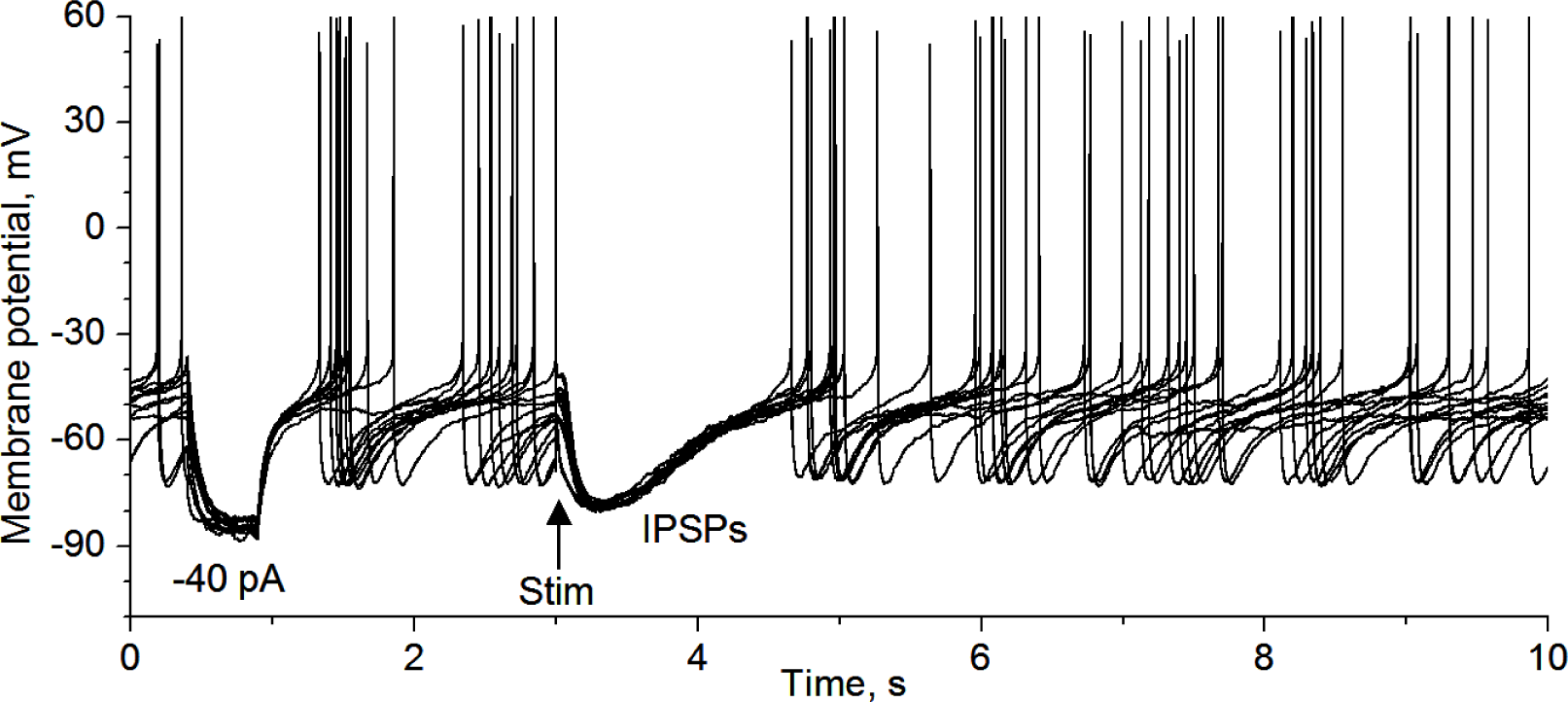
IPSPs inhibit raphe 5-HT neuron firing. KSO_3_CH_4_-based intracellular solution. The first hyperpolarization pulse was induced by −40 pA pulse to monitor input resistance and to provide an estimate of the strength of synaptic input underlying the IPSPs: for this 5-HT neurons, the synaptic current was less than 40 pA. 10 sweeps are displayed. Stim, local electrical stimulation.

### 5-HT inhibition of GABAergic inputs to DR 5-HT neurons

In the presence of 10 μM CNQX and 20 μM D-APV to block fast ionotropic glutamate receptors, we recorded local extracellular stimulation-evoked inhibitory postsynaptic currents (evoked IPSC or eIPSC) in 5-HT neurons voltage-clamped at −70 mV; to observe the repetitive GABA release properties of the afferent axon terminals, we used a train stimulation with an intra-train stimulation frequency of 10 Hz, a common spiking frequency of the neurons projecting to the raphe. At this frequency, the axon terminal can apparently sustain the IPSC amplitude or GABA release--after a moderate depression during the first several IPSCs (**Fig. 3A**). These results indicate that GABA afferents can effectively and faithfully inhibit DRN 5-HT neurons, according to activity of the presynaptic afferent GABA neurons.

**Fig. 3.**
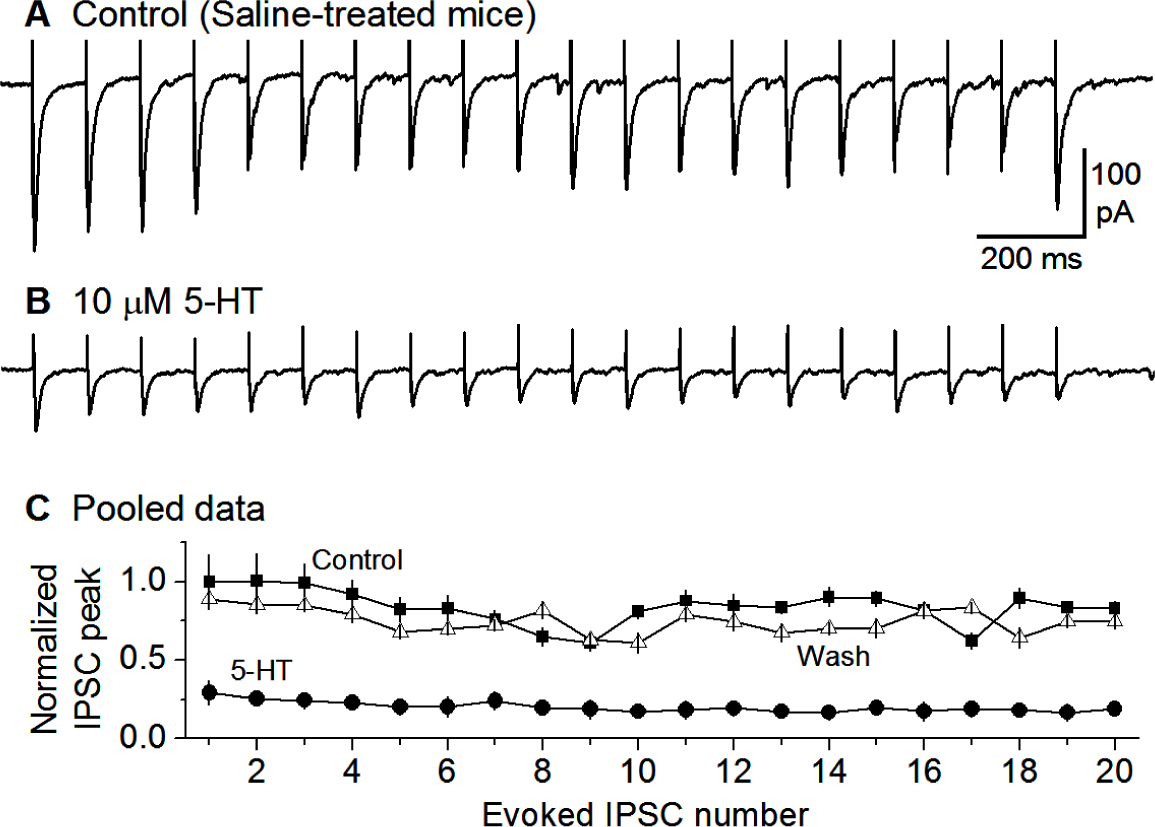
Bath-applied 5-HT substantially reduced the eIPSCs recorded in DRN 5-HT neurons in salinetreated control mice. A train of 20 10-Hz stimuli evoked 20 IPSCs under a basal condition (**A**) and during application 10 µM 5-HT (**B**). For comparison, each IPSC peak was normalized to the first IPSC peak under control condition (**C**).

To examine the potential effects of 5-HT on GABAergic inputs to raphe 5-HT neurons in normal mice, we electrically evoked a train of 20 repetitive eIPSCs using a train of 20 stimuli with an intra-train frequency of 10 Hz. After a stable baseline recording, bath application of 10 μM 5-HT produced a reversible reduction of the peak amplitude of each eIPSC in the train by about 70% (n=6 neurons; **Fig. 3B,C**); also noteworthy is that the peak amplitude of each IPSC in the train was reduced by a similar magnitude such that the paired pulse ratio (PPR) was changed only minimally.

### Girk channel blocker tertiapin-Q prevents 5-HT inhibition of GABAergic inputs to 5-HT neurons

Our observation that 5-HT reduced the amplitude of repetitive eIPSCs globally without changing PPR suggested that 5-HT, probably via 5-HT1B receptor activation, may reduce the general excitability and hence transmitter release of the GABAergic axon terminals synapsing onto the 5-HT neurons. One possibility or possible mechanism is that 5-HT activates Girk channels expressed in the GABAergic afferent axon terminals—recent anatomical and functional studies support this possibility in both the raphe and other brain areas (Fernández-Alacid et al. 2009, 2011; Ladera et al. 2008; Llamosas et al. 2017). In raphe 5-HT neurons, somatodendritic 5-HT1A couples to Girk channels that causes hyperpolarization and hence inhibition of spike firing, accompanied by decreased whole-cell input resistance and excitability. We hypothesized that Girk channels expressed on raphe afferent axon terminals may reduce axonal excitability, leading to a general decrease in membrane excitability, fewer spikes (some axon terminals fail to spike), less Ca influx, and hence a general decrease in GABA release.

To test this hypothesis, we used tertiapin-Q that is a known selective inhibitor of Girk channels (Chen and Johnston 2005; Kim and Johnston 2015; Montalbano et al. 2015; Llamosas et al. 2017). As shown in **Fig. 4A-D**, in the presence of 1 μM tertiapin-Q, bath application of 10 μM 5-HT did not reduce eIPSCs (n = 6). These results indicate that blockade of GIRK channels is a key mechanism for 5-HT to inhibit the GABAergic inputs to and GABA_A_R IPSCs in raphe 5-HT neurons.

**Fig. 4.**
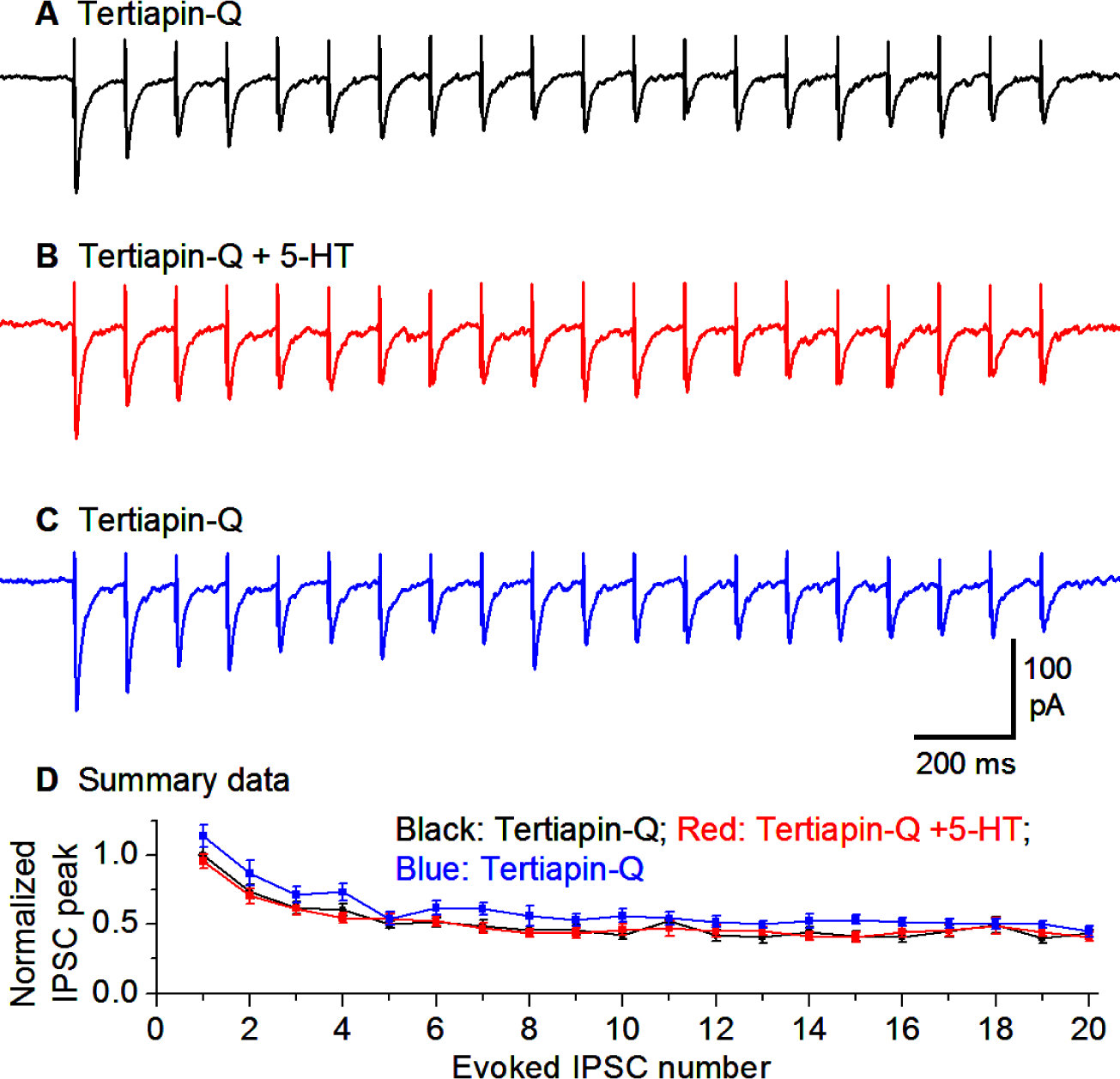
Girk channel blocker tertiapin-Q prevents 5-HT inhibition of IPSCs. A train of 20 10-Hz stimuli evoked 20 IPSCs in the presence of tertiapin-Q (**A**) and tertiapin-Q and 10 µM 5-HT (**B**) and after washing out 5-HT (**C**). For comparison, each IPSC peak was normalized to the first IPSC peak under control condition (**D**).

### Chronic fluoxetine treatment down-regulates 5-HT inhibition of GABA inputs to 5-HT neurons

Chronic fluoxetine treatment is known to down-regulate or desensitize 5-HT1A autoreceptors on the somatodendritic areas and axon terminals of raphe 5-HT neurons (via decreased receptor expression, receptor-G_i/o_ coupling, and/or down-regulation of downstream signaling pathway). A similar down-regulation for 5-HT1B receptors on non-5-HT neuron axon terminals is entirely possible but has not been reported. We hypothesized that chronic fluoxetine may induce a desensitization or downregulation of presynaptic 5-HT1B inhibition of IPSCs in raphe 5-HT neurons.

To test this hypothesis, we treated the mice for ≥ 14 days with fluoxetine (10 mg/kg per day) with saline injection as control. Then we examined the effects of bath-applied 5-HT (10 μM) on the GABAergic IPSCs in raphe 5-HT neurons in fluoxetine-treated mice. We found that 10 μM 5-HT decreased 10 Hz stimulus-train-evoked eIPSC amplitude only minimally (n=9) (**Fig. 5A-D**). These results indicate that chronic treatment with fluoxetine down-regulated or reduced the 5-HT inhibition of GABA inputs to 5-HT neurons.

**Fig. 5.**
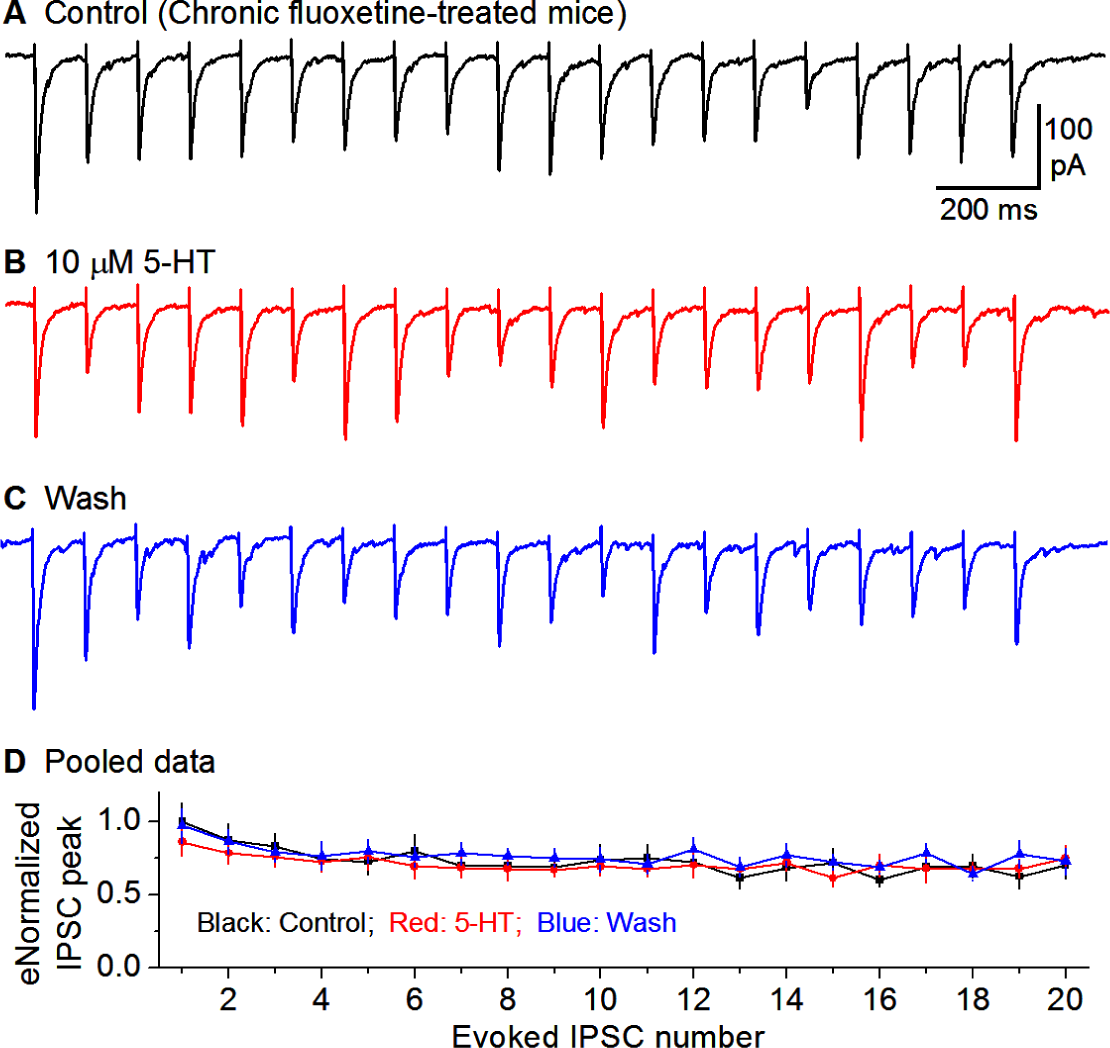
Chronic fluoxetine treatment downregulates 5-HT inhibition of GABA inputs to 5-HT neurons. Recorded with KCl-based intracellular solution. A train of 20 10-Hz stimuli evoked 20 IPSCs under basal condition (**A**), during 10 µM 5-HT application (**B**) and after washing out 5-HT (**C**). For comparison, each IPSC peak was normalized to the first IPSC peak under control condition (**D**).

### Chronic fluoxetine treatment down-regulates 5-HT neuron autoinhibition

5-HT, by activating somatodendritic 5-HT1ARs in 5-HT neurons, activates Girk channels in 5-HT neurons, leading to hyperpolarization and inhibition of these neurons (Bayliss et al. 1997). Chronic fluoxetine treatment has been shown to downregulate 5-HT1A receptors (Blier and El Manari 2013; Descarries and Riad 2013; Hensler 2002) and remove the initial 5-HT neuron firing inhibition after SSRI administration (Blier and de Montigny 1983; Czachura and Rasmussen 2000). Here, we used whole-cell recording to directly determine if chronic fluoxetine treatment reduces the 5-HT-activated Girk current-induced hyperpolarization and the associated low excitability in raphe 5-HT neurons.

In saline-treated mice, bath application of 10 μM 5-HT induced a robust hyperpolarization in raphe 5-HT neurons (−26.03±1.82 mV, n=8; **Fig. 6A1-A3, C1**); this hyperpolarization was accompanied by a substantial decrease in input resistance (584.05±23.14 MΩ under control, 128.17±5.65 MΩ under 5-HT n=8; **Fig. 6A1-A3, C2**); further, current injection-evoked spike firing was severely inhibited (10.7±0.6 spikes under control, 0 spike under 5-HT, n=8; **Fig. 6A1-3**). These results are fully consistent with 5-HT activating 5-HT1ARs and opening Girk channels.

**Fig. 6.**
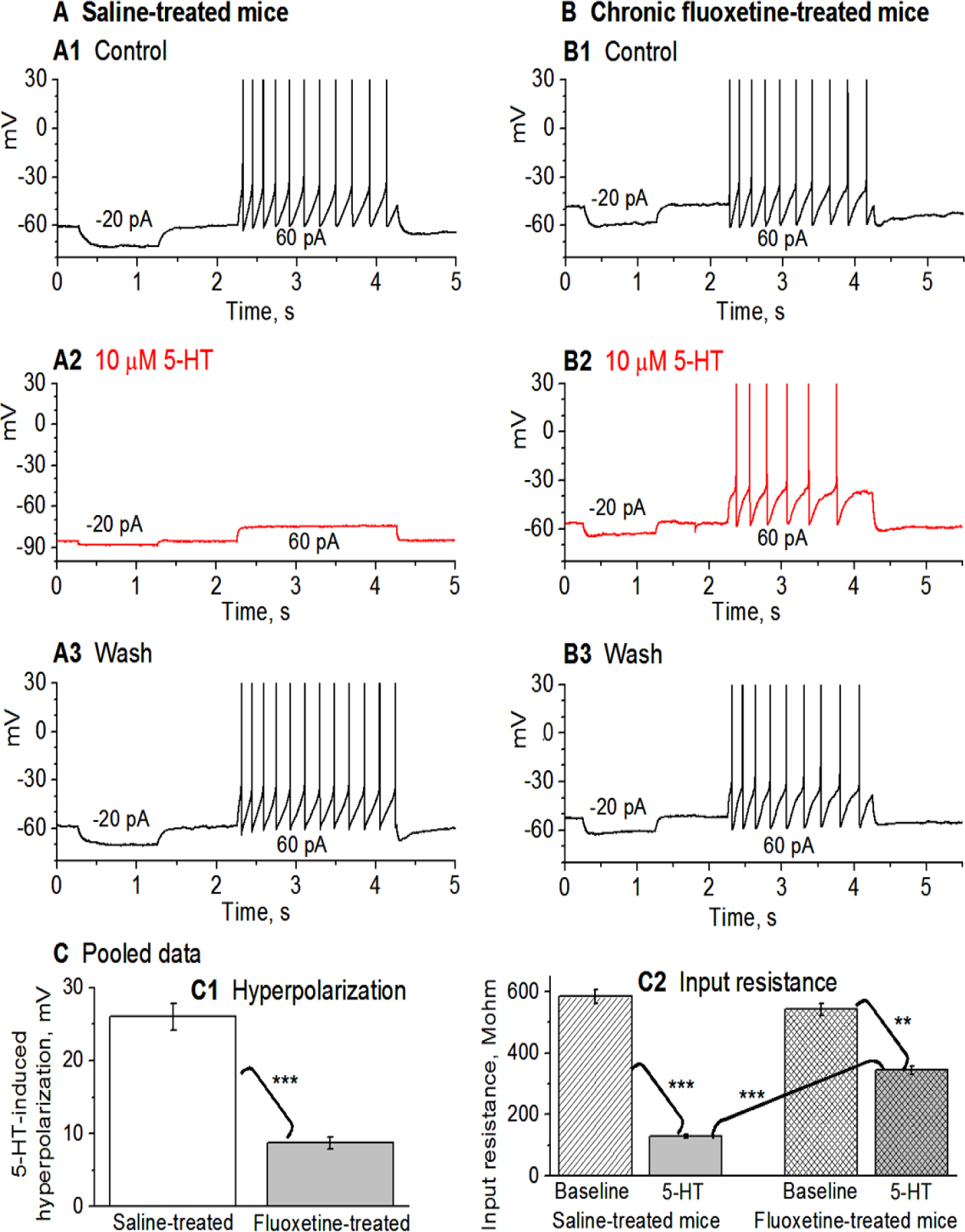
Chronic fluoxetine treatment renders DRN 5-HT neurons resistant to 5-HT autoinhibition by substantially downregulating 5-HT autoinhibition of DR 5-HT neurons. **A (A1-A3)**: In brain slices from saline-treated control mice, bath-application of 10 µM 5-HT caused a large (26 mV) hyperpolarization, a large decrease in input resistance and a cessation of the evoked spike firing. **B (B1-B3)**: In brain slices from fluoxetine-treated mice, bath-application of 10 µM 5-HT caused a much smaller (10 mV) hyperpolarization, a smaller decrease in input resistance and a modest decrease of the evoked spike firing. **C (C1, C2)**: Summary of the data. ***, p<0.001; **, p<0.01.

In mice treated with fluoxetine for 14 days, the responses to 5-HT were much smaller. Under an identical recording condition, bath application of 10 μM 5-HT induced a much smaller hyperpolarization in raphe 5-HT neurons (−8.7±0.9 mV, n=8; **Fig. 6B1-B3, C1**); this smaller hyperpolarization was accompanied by a smaller decrease in input resistance (542.04±20.20 MΩ under control, 343.91±14.26 MΩ under 5-HT, n=8; **Fig. 6B1-B3, C2**); further, current injection-evoked spike firing was only moderately reduced (10.5±0.5 spikes under control, 7.5±0.5 spike under 5-HT, n=8; **Fig. 6B1-B3**). These results indicate that in chronically fluoxetine-treated mice, 5-HT/5-HT1ARs-induced Girk current in raphe 5-HT neurons is reduced. We also need to note here that although 10 μM 5-HT still inhibited DRN 5-HT neurons in brain slices from chronic fluoxetine-treated mice, the endogenous extracellular 5-HT level is more likely around 1 μM that probably still can substantially inhibit 5-HT neurons in untreated or saline-treated animals, but cannot inhibit these 5-HT neurons in SSRI-treated animals. These experiments to determine the dose-response curves need to be performed in future studies.

## Discussion

The main findings of our present study are that (1) 5-HT, probably via 5-HT1B activating presynaptic Girk channels, reduces GABAergic regulatory inputs to raphe 5-HT neurons, and (2) chronic treatment with the antidepressant fluoxetine downregulates this 5-HT inhibition of GABAergic inputs and also the 5-HT-activated somatodendritic Girk current-mediated hyperpolarization and autoinhibition. Below, we will discuss these results and also their weaknesses.

### Chronic fluoxetine treatment enhances GABAergic inhibitory influence on dorsal raphe 5-HT neurons by downregulating presynaptic 5-HT inhibition

In this study, we first demonstrated that IPSPs inhibit the spike firing of DRN 5-HT neurons, probably due to their high input resistance such that even a small inhibitory synaptic input can cause a significant hyperpolarization and hence an inhibition of spike activity. These results are new and useful and expand prior studies documented that DRN 5-HT receive inhibitory synaptic inputs but did not show explicitly that these IPSPs inhibit spike firing in DRN 5-HT neurons (Lemos et al. 2006; Morikawa et al. 2000; Pan and Williams 1989; Williams et al. 1988).

Second, our data indicate that 5-HT reduced the GABAergic inputs to DRN 5-HT neurons, probably by activating presynaptic 5-HT1B receptors on GABA afferent terminals synapsing on 5-HT neurons based on the well established observation that 5-HT1B receptors are commonly on axon terminals (Boschert et al. 1994; Ding et al. 2013, 2015; Lemos et al. 2006; Li and Bayliss 1998; Morikawa et al. 2000; Sari 2014), although in the present study we did not use 5-HT1B receptor-selective ligands to test this possibility. This mechanism serves to reduce the inhibitory influence of local GABA neurons and external GABAergic centers such as the hypothalamus, lateral habenula nucleus, RMTg, lateral preoptic area and the pontine ventral periaqueductal gray and VTA, SNr have on DRN 5-HT neurons (Gervasoni et al. 2000; Kirouac et al. 2004; Lavezzi et al. 2012; Pollak et al. 2014; Reisine et al. 1982; Sego et al. 2014; Soiza-Reilly and Commons 2014; Taylor et al. 2014; Zhou et al. 2017). This mechanism may contribute to the overall regulation of 5-HT neuron spiking activity that matches the animal’s mental and behavioral status and needs.

Third, our data indicate that pretreatment with the Girk channel inhibitor tertiapin-Q prevented the inhibitory effect of 5-HT on the IPSCs. One interpretation is the following. 5-HT activates presynaptic 5-HT1 receptors (5-HT1A and/or 5-HT1B) that in turn activate Girk channels expressed at the axon terminals, thus reducing axon terminal excitability, leading to fewer spikes, less Ca influx and less GABA release. Girk channels have been reported to express at axon terminals in multiple brain areas and also at afferent axon terminals in DRN (Fernández-Alacid et al. 2009, 2011; Ladera et al. 2008; Llamosas et al. 2017; Ponce et al. 1996). This is also consistent with the report that presynaptic DA receptors reduce GABA release onto midbrain DA neurons by activating presynaptic Girk channels (Michaeli and Yaka 2010). However, we recognize that the postsynaptic 5-HT1A receptor-activated Girk channels may be a confounding factor, although it has been reported that 5-HT1A agonism did not alter the mIPSC amplitude in DRN 5-HT neurons recorded with K-based intracellular solution (Lemos et al. 2006), arguing against the possibility of lower input resistance contributing to lower eIPSC amplitude. To fully resolve this issue, future studies need to use CsCl-based intracellular solution to block postsynaptic Girk channels. Also, since our current study did not definitively identify the 5-HT receptors involved, future studies will need to use selective 5-HT1A ligands and 5-HT1B ligands together with transgenic 5-HT1A knockout mice and 5-HT1B knockout mice to identify the 5-HT1 receptor subtype mediating the 5-HT effects observed here.

Finally, we found that this presynaptic 5-HT inhibition was downregulated after 2-week chronic fluoxetine treatment. Previous studies have established chronic antidepressant treatment desensitizes 5-HT1 autoreceptors on 5-HT axon terminals inhibiting 5-HT release, and this downregulation temporally coincided with the onset of antidepressant effects in the animal models (Anthony et al. 2000; Blier and El Mansari 2013; Neumaier et al. 1996; Newman et al. 2004; Tiger and Lundberg 2018). Our present study suggests that chronic SSRI treatment and hence extracellular 5-HT increase may affect 5-HT1B heteroreceptors. Thus, it appears that chronic exposure to high levels of extracellular 5-HT can desensitize the function and/or decrease the cell surface or de novo expression of both 5-HT1B autoreceptors and heteroreceptors. The reduced 5-HT inhibition indicates that after chronic fluoxetine treatment, 5-HT neurons can be more effectively influenced by outside GABAergic neurons in the hypothalamus, substantia nigra pars reticulata, ventral tegmental area, and other brain areas including the cerebral cortex and brainstem (Soiza-Reilly and Commons et al. 2014).

We also need to note here that literature data indicate that 5-HT1B receptors are probably expressed in GABA neurons in the raphe and on the axon collaterals of raphe 5-HT neurons; these 5-HT1B receptors can likely inhibit GABA and 5-HT release, but their functional roles in normal animal physiology and depression pathogenesis are being investigated but not established (Lemos et al. 2006; McDevitt et al. 2011; McDevitt RA, Neumaier 2011; Tiger et al. 2011).

### Chronic antidepressant treatment renders DRN 5-HT neurons resistant to 5-HT autoinhibition by downregulating 5-HT inhibition of the intrinsic excitability

We found that 2-week daily fluoxetine treatment substantially reduced 5-HT-induced autoinhibition of DRN 5-HT neurons monitored by whole-cell patch clamp recording. Our present results are consistent with and expand the literature data in the field. In their pioneering study using in vivo extracellular spike recording, Blier and De Montigny (1983) found that ip injection of the SSRI zimelidine initially autoinhibited the extracellularly recorded spontaneous spike firing in DRN 5-HT in rats; however, following repeated daily treatment with this SSRI, the autoinhibition gradually declined and eventually disappeared after 2 weeks of treatment, demonstrating that chronic SSRI treatment that induced an chronic increase of extracellular 5-HT level can desensitize or downregulate 5-HT autoinhibition. In a more detailed study with 3 doses of fluoxetine (5, 10 and 20 mg/kg per day) combining extracellular spike recording, Czachura and Rasmussen (2000) confirmed that fluoxetine-induced DRN 5-HT neuron autoinhibition, pronounced within the first 3 days of fluoxetine treatment, disappeared after 14-21 days of daily fluoxetine treatment.

Our present study provides complementary whole-cell recording data that add new and useful information about the changes in input resistance, intrinsic excitability and Girk current during 5-HT stimulation after chronic fluoxetine treatment--these important details could not be recorded in prior in vivo studies recording extracellular spikes; specifically, our data show that after 2 weeks of daily IP 10 mg/kg fluoxetine treatment, DRN 5-HT neurons became more resistant to 5-HT autoinhibition by producing a much smaller Kir current, a much smaller hyperpolarization, a much smaller decrease in input resistance--these new whole-cell data are consistent with and solidify prior extracellular spike data; hence in the presence of fluoxetine or another SSRI type antidepressant, 5-HT neurons can maintain the spiking activity (∼ 1 Hz) and release 5-HT in the projection areas while the SSRI blocks the reuptake and increases extracellular 5-HT level, the intended pharmacological effect that is believed to underlie antidepressant therapeutic efficacy.

Literature evidence indicates that the following chain of events may underlie the chronic fluoxetine-induced downregulation of 5-HT neuron autoinhibition. Fluoxetine blocks SERT-mediated 5-HT reuptake and thus increase the extracellular 5-HT level. Chronic increase in the extracellular 5-HT level can desensitize the 5-HT1A and 5-HT1B inhibitory autoreceptors in 5-HT neuron cell bodies and their axon terminals via diminished receptor-G-protein coupling (Castro et al. 2003; Hensler 2002; Cornelisse et al. 2007; Li et al. 1996, 1997; Newman et al. 2004), receptor internalization (Descarries and Riad 2012), or reduction of de novo receptor expression (Anthony et al. 2000; Neumaier et al. 1996), thus removing autoinhibition and leading to sustained increase in extracellular 5-HT and thus contributing to the antidepressant effect (Blier and El Mansari 2013; Wong et al. 2005). The situation with presynaptic 5-HT1B receptors on non-5-HT neurons after SSRI treatment is understudied and less clear. Our present study suggests that the downregulating mechanisms may be similar to those of 5-HT1A and 5-HT1B autoreceptors.

### Functional implications

Our present finding that chronic fluoxetine downregulates 5-HT inhibition of GABAergic synaptic inputs to DRN 5-HT neurons suggest that chronic antidepressant treatment can enable extrinsic, behaviorally important GABA neuron activity to more effectively and closely influence DRN 5-HT neurons such that 5-HT neuron activity and hence 5-HT release better match behavioral needs. Equally important, our second finding that chronic fluoxetine treatment reduces somatic 5-HT autoreceptor (likely 5-HT1A receptor)-activated Girk channel-mediated hyperpolarization and decrease in input resistance and intrinsic excitability complement the results of prior in vivo extracellular spike recording studies, and indicate that chronic antidepressant treatment can render DRN 5-HT neurons resistant to 5-HT autoinhibition and leading to increased 5-HT neuron activity and 5-HT release. These cellular events are likely key pharmacological mechanisms by which SSRIs exert their antidepressant therapeutic effects.

## Acknowledgements

This work was supported by NIH grant R01NS097671. Wei Zhang was a recipient of an award from China Scholarship Council.

